# Bumblebees do not prefer consistent floral scents over variable ones

**DOI:** 10.1101/2025.03.15.643418

**Authors:** Mélissa Armand, Lisa Zeilmann, Christian Weinzettl, Tomer J. Czaczkes

## Abstract

To attract pollinators, flowering plants evolve diverse sensory traits into compelling signals. Floral scent, in particular, plays a key role in drawing bees from a distance and shaping their foraging choices. Scent composition varies widely, including across flowers of the same plant species. Yet, it is unclear whether scent variability influences bee flower choices, and thus whether plants would be under selection to minimise variation in their scent composition. Since bees typically avoid variability in rewards, we hypothesised they would favour flowers with more consistent scents. To test this, we trained individual bumblebees (*Bombus terrestris*) on two equally rewarding flower arrays: one with a consistent scent blend across flowers, and the other with variable scent blends between flowers. Contrary to expectations, bees showed no preference for scent consistency. They readily foraged from both arrays across bouts and did not favour either flower type in the binary choice test. To the best of our knowledge, this is the first study to examine how bees respond to scent variability. A better understanding of scent profile preferences in pollinators could offer new insights into their co-evolution with plants and the development of floral traits. More broadly, further research is needed on how pollinators respond to variability and unpredictability in neutral cues like scent or colour, a largely overlooked aspect of foraging decision-making.

## INTRODUCTION

Most flowering plants depend on pollinators for reproduction (Klein et al., 2007), so their flowers act like billboards, using various colours, shapes, and scents to attract them (Dobson, 1994). Floral scents, especially, play a key role in pollination (Burkle et al., 2020): scent bouquets attract bees from far away (Raguso, 2004) and often serve as bees’ primary cue for deciding whether to land on a flower (Kunze et al., 2001; Raguso, 2008a; Sprayberry, 2018). Yet, despite their influence on bees’ foraging decisions (Farré-Armengol et al., 2015; Larue et al., 2016), the role of scent in flower choices remains understudied in comparison to flower colours and shapes (Fenster et al., 2004; Raguso, 2008b).

Floral scents consist of complex blends of volatile organic compounds (Knudsen et al., 2006), released from various parts of the flower (Pichersky et al., 1994; Raguso and Pichersky, 1999). These scent bouquets act as unique floral identifiers (Raguso, 2008a) and are highly diverse across plant species (Raguso and Pichersky, 1999; Levin et al., 2003). Scent composition can also vary across plant populations and even among individual flowers within the same species (Raguso et al., 2003; Burdon et al., 2015; Delle-Vedove et al., 2017). However, the causes of intraspecific scent variation remain poorly studied and understood (Majetic et al., 2009; Raguso, 2020), as well as its effects on pollinator foraging behaviour.

Scent cues allow flower-visiting insects to associate flowers with pollen and nectar rewards (Kevan and Baker, 1983; Wells and Wells, 1985; Giurfa, 2007), and help them distinguish between plant species (Dobson, 1994). Both honeybees and bumblebees can detect differences in floral scent blends (Laloi and Pham-Delègue, 2004; Wright et al., 2005), and small variations in compound ratios were shown to strongly affect their flower choices (Dobson, 1994; Raguso, 2004; Tan and Nishida, 2012; Solís-Montero et al., 2018). Like floral colours, bees have innate preferences for certain floral volatiles or scent blends (Raguso, 2008a; Schiestl and Dötterl, 2012). However, it is unclear whether bees also prefer certain patterns of scent variability.

Floral scent is a highly variable trait that plants can adjust to reflect their current state (Dudareva et al., 1996; Dobson, 1994). For instance, scented nectar can inform pollinators about reward availability (Heinrich, 1979; Raguso, 2004; Gervasi and Schiestl, 2017). Plants can also modify their scents to better attract pollinators (Dudareva et al., 2004; Raguso, 2008a; Leonard et al., 2011). For example, some flowers adjust the intensity of their scent emissions throughout the day to match pollinator activity (Loughrin et al., 1990; Raguso, 2008b; Wright and Thomson, 2005). Scent composition may also vary at different stages of flower development (Majetic et al., 2015; Burkle et al., 2020). Do bees favour plant species with more consistent or variable scent profiles across flowers?

Pollinators not only respond to floral traits but also influence how these traits evolve through selection pressure, including scent (Parachnowitsch et al., 2013; Schiestl and Johnson, 2013; Ollerton et al., 2011). Stabilising selection driven by pollinator preferences is well-established for flower colour (Goulson, 1999; Whibley et al., 2006), and similar selection may also shape scent composition (Huber et al., 2005; Mant et al., 2005; Salzmann et al., 2007). The pollination syndrome hypothesis suggests that unrelated plant species visited by similar pollinators tend to develop similar floral traits, such as scent composition (Fenster et al., 2004; Dobson, 2006; Farré-Armengol et al., 2020), although this idea remains debated (Rosas-Guerrero et al., 2014; Ollerton et al., 2015).

Bumblebees tend to specialise and forage on a few flower species (Heinrich, 1979; Chittka et al., 1999). Such flower constancy benefits plants by ensuring that bees repeatedly visit the same species, increasing pollination efficiency (Heinrich, 1977; Waser, 1986). Like colours, scents promote flower constancy in pollinators (Gegear and Laverty, 2005; Gegear, 2005). In turn, this flower constancy probably contributes to stabilising floral traits, including scent profiles (Chittka et al., 1999; Goulson, 1999). This tendency to be flower constant may reflect a broader preference for consistency in bee foraging decisions. In fact, bees typically exhibit an aversion to variability and unpredictability in rewards (see review by Anselme, 2018). Similarly, we propose that bees may favour plant species with more consistent scent profiles across flowers, potentially placing selection pressure on plants to tightly control their scent composition.

Here, we investigated whether bumblebees (*Bombus terrestris*) prefer flowers with either consistent or variable scents. To test this, we trained individual bees on two equally rewarding arrays of artificial flowers: all yellow or all green. In one array, all flowers contained sucrose solution scented with a fixed ratio of two artificial food flavourings, creating a consistent scent bouquet. In the other array, flowers had varying ratios of a different pair of scents, making varying, unpredictable scent bouquets. We hypothesized that bees would favour the consistent scent flower, being more predictable than the other flower. We predicted that in a binary choice test, bees would first visit the flower colour associated with the consistent scent.

## MATERIAL AND METHODS

### Colony setup

Commercial *Bombus terrestris* colonies were purchased from Koppert (The Netherlands) and kept under controlled laboratory conditions at 22–24°C with a 14:10 light:dark cycle. Colonies were housed in wooden nestboxes, each connected to its respective flight arena (60 × 50 × 35 cm) leading to a small chamber (6 × 5 × 3 cm). The chamber featured a second tube that provided direct access to the arena and was fitted with transparent, removable shutters to regulate bee movement between the nest and arena (see apparatus **Fig. 1**).

**Figure 1:**
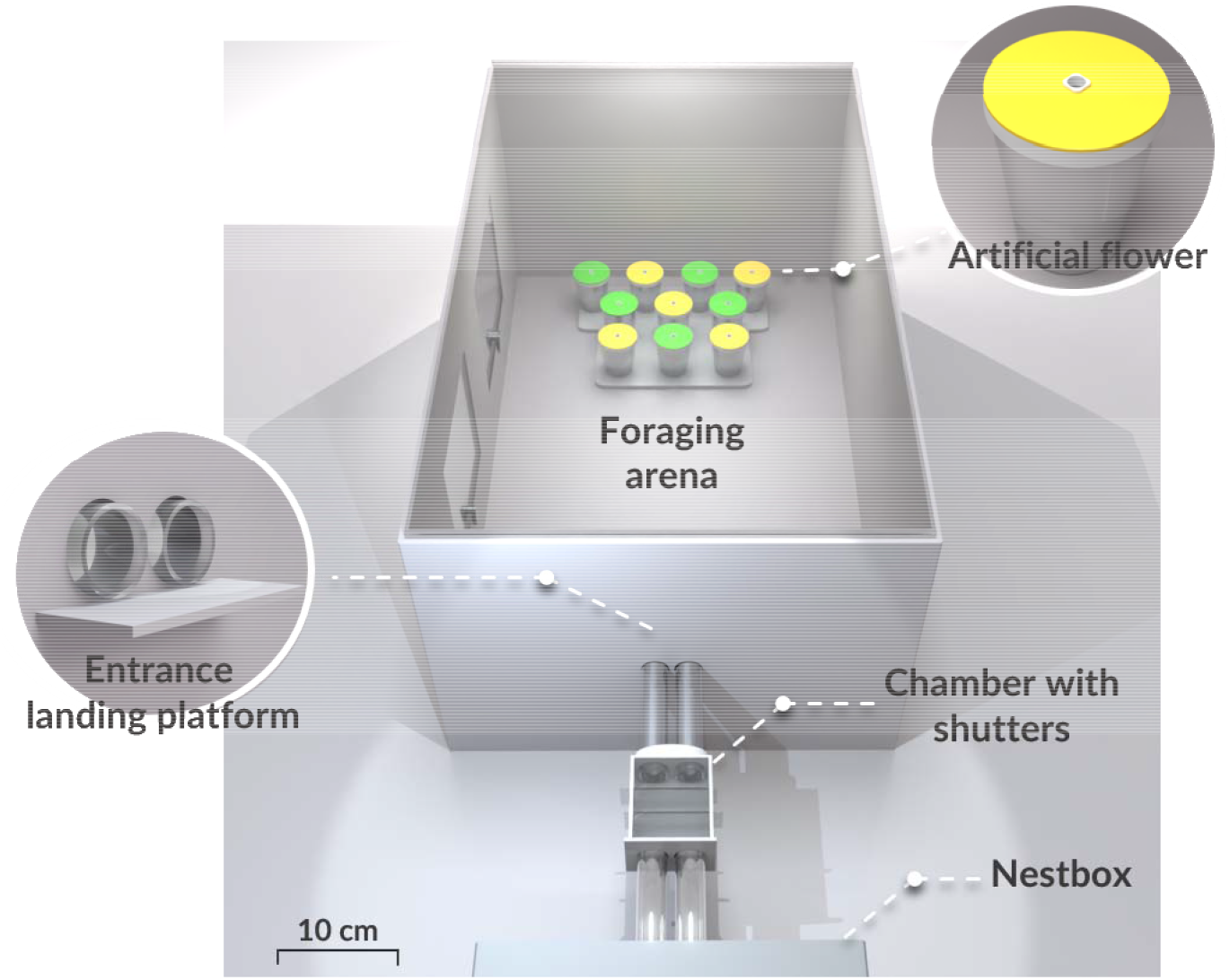
Top view of the experimental setup, to scale. The 3D model, created in Blender, illustrates a binary choice bout with a mixed flower array of yellow and green flowers, arranged in a “yellow biased” layout (*i*.*e*., two yellow flowers on the front row).

Bees were provided daily with pollen balls made from a mix of organic flower pollen pellets and 35% (w/w) sucrose solution, placed directly in the nestboxes. During the day, workers foraged freely on artificial flowers in the flight arena, which offered 35% (w/w) sucrose solution and were regularly refilled. Active foragers were captured and marked on the thorax with uniquely numbered, coloured tags, and considered for selection in the experiment on the same day. A total of 48 bees participated in the experiment from five colonies in May–June 2022 (see colony details in Supplement **S1**).

### Artificial flowers

Each artificial flower consisted of a transparent plastic cup with a white lid (height: 4.5 cm, diameter: 3.8 cm), topped with a 2 mm-thick disc of coloured rubber foam. At the centre, a small opaque white resin cup (diameter: 4 mm, depth: 6 mm) was inserted into a pre-cut hole and filled with sucrose solution (**Fig. 1**).

In the experiment, flowers were either yellow (peak reflectance: ∼520–700 nm) or green (peak reflectance: ∼530 nm; see **Supplement S2** for reflectance curves). We selected these colours for their perceptual similarity while still allowing bees to differentiate them (see control experiment below), and to avoid strong innate preferences typically observed for blue or violet flowers (Gumbert, 2000; Raine and Chittka, 2007). Between experimental sessions, bees were pre-trained on bicoloured, half green half yellow flowers, to ensure they associated sucrose rewards equally with both flower colors used in the experiment (Raine and Chittka, 2008).

During training, bees foraged on an array of flowers consisting of a 3 × 3 grid of either all-yellow or all-green flowers. In the binary choice test, the array contained 5 green and 5 yellow flowers, arranged in two front rows of three flowers and a third row of four flowers (see **Fig. 1** below). Flowers were spaced 3.5 cm apart and mounted on a grey-painted plate matching the all-grey arena. The plate ensured consistent flower placement and allowed for easy flower replacement between foraging bouts.

### Scented sucrose solutions

Artificial flowers provided 35% w/w sucrose solution at a fixed volume, determined individually for each bee based on an estimate of its honey crop size (see below). Sucrose solutions were scented using strawberry, rose, lemon, or vanilla food flavourings (Seeger, Springe, Germany). We created six artificial scent bouquets by mixing the flavourings in different combinations: strawberry–rose, lemon– vanilla, strawberry–vanilla, lemon–rose, strawberry–lemon, and vanilla–rose. Each combination was prepared in three different ratios: 1:1 (equal parts of both scents), 1:3 (25% scent A, 75% scent B), or 3:1 (75% scent A, 25% scent B).

Bees were familiarised with each scent (strawberry, rose, lemon and vanilla) 48 hours before the experiment, by placing filter papers soaked with 25 μL of each food flavouring in the four corners of the nestboxes, allowing airborne scents to disperse. This ensured that bees were not reluctant to collect sucrose solution during the experiment due to unfamiliar scents.

### Estimation of crop size

To determine the appropriate volume of sucrose solution used per flower in the experiment, we first estimated the crop capacity of individual bees. Bee foraged on a 3 × 3 array of bicoloured flowers (half green, half yellow), with each flower providing 15 μL of a 35% w/w sucrose solution. We recorded the number of flowers collected over two consecutive foraging bouts and estimated the crop size of the bees by averaging the total volume collected across both bouts. This average volume was then divided by nine, and the resulting volume was used as the reward per flower during the experiment. This ensured that the bees could collect all available rewards in each foraging bout.

### Training

After its crop size was estimated, each bee was trained for six consecutive foraging bouts on two different 3 × 3 flower arrays, alternating between them in each bout. Each flower array was associated with a flower colour (green or yellow) and a scent combination (strawberry–rose, lemon– vanilla, strawberry–vanilla, lemon–rose, strawberry–lemon, or vanilla–rose).

Bees experienced a distinct scent combination for each flower colour; for example, if one array had green flowers scented with strawberry–lemon, the other had yellow flowers scented with the remaining two scents (e.g., rose–vanilla). We assigned different scent combinations to each flower colour to ensure clear differentiation between flower types and allowed the use of three different scent ratios (1:1, 1:3, and 3:1) in the variable flower type, while maintaining a consistent 1:1 ratio in the consistent flower (see below).

The flower arrays were one of two types:

1. **Consistent flower array**: All flowers in the array had the same scent combination at a fixed 1:1 ratio (equal parts of both scents);
2. **Variable flower array**: Flowers in the array had varying scent ratios: some 1:1, some 1:3 (25% scent A, 75% scent B), and some 3:1 (75% scent A, 25% scent B).

For example, if a bee experienced green flowers as the consistent array, all green flowers would have strawberry–lemon in a fixed 1:1 ratio, and the yellow flowers in the variable array would contain rose– vanilla with varying scent ratios (1:1, 1:3, and 3:1).

Half of the bees experienced the consistent flower array as green and the variable array as yellow (n = 24 bees), and the other half the reverse (n = 24 bees). Bees were required to consistently collect sucrose solution from both array types to ensure they perceived each scent combination as an acceptable reward. Flowers were replaced with clean ones each new bout to prevent scent marks from influencing subsequent foraging (Goulson et al., 2000; Saleh et al., 2007).

### Binary choice test

After the six training bouts, each bee was presented with an array of 10 flowers — 5 yellow and 5 green, arranged so that neighbouring flowers alternated in colour. The flowers were unrewarded and filled with unscented plain water. Bees were randomly assigned one of two array layouts, where the front row contained either two green flowers and one yellow (“green-biased” layout), or the reverse (“yellow-biased” layout). We recorded the first flower choices as an indicator of preference.

### Control experiment

A pilot was conducted before the main experiment to test whether bees could differentiate between green and yellow flowers. The setup mirrored the main experiment, where individual bees completed six training bouts to a 3 × 3 flower array of a single colour and alternating between arrays each bout, followed by a binary choice test with both flower colours.

Instead of being scented, one flower type offered 25 μL of a low-quality sucrose solution (15% w/w), while the other provided 35% w/w sucrose solution. This clear difference in reward quality allowed us to assess whether bees could correctly identify the flower colour associated with the higher-quality reward. All tested bees (n = 10) successfully chose first the flower colour associated with high-quality reward in the final test.

### Data analysis

The binary choice test was video-recorded for each bee with a camera (Sony HDR-CX220) positioned above the flight arena. Bee behaviour was analysed using the event-logging software BORIS (v8.6). We tested the hypothesis that bees would favour the flower colour (green or yellow) associated with a consistent scent. To assess this, we recorded (1) the first flower visit and (2) the first 10 flower visits during the binary choice test, as indicators of preference.

1. For the first flower visit, we built a generalized linear mixed model (GLMM) with a binomial distribution, using first flower scent choice (1 for consistent, 0 for variable) as the response variable and flower colour (green or yellow) as the predictor. Random effects included bee colony, first flower scent encountered (i.e., whether the first training bout involved the consistent or variable array), first flower colour encountered (green or yellow in the first training bout), and binary test layout (“green-biased” or “yellow-biased”). Predicted probabilities of first choice were assessed through post hoc pairwise comparisons with a Tukey correction.
2. For the first ten flower visits, we fitted a GLMM with a binomial distribution, using flower scent choices (1 for consistent, 0 for variable) as the response variable and flower colour (green or yellow) as the predictor. Random effects included individual bee identities nested within their colony, first flower scent encountered, first flower colour encountered and binary test layout. Predicted probabilities of flower choices were evaluated through post hoc pairwise comparisons with a Tukey correction.

Data processing was conducted in Python (v3.11, Python Software Foundation, 2023) using the *pandas* library (McKinney, 2010) for data structuring and *seaborn* (Waskom, 2021) and *Matplotlib* (Hunter, 2007) for data visualization. Statistical analyses were performed in R (v4.1, R Core Team, 2022) using the *glmmTMB* package (Brooks et al., 2017) for GLMMs, and *emmeans* (Lenth, 2020) for post hoc tests. Model residuals were evaluated with the *DHARMa* package (Hartig, 2020). Complete statistical analyses and datasets are available on Zenodo (https://doi.org/10.5281/zenodo.14993082).

## RESULTS

### First flower choice

31 of the 48 tested bees (64.6% ± 8.6%) first visited a yellow flower in the binary choice test, while 17 bees (35.4% ± 11.6%) chose a green flower. This difference was borderline significant (*p* = 0.059, two-tailed exact binomial test, 95% CI: 49.5–77.8%). Regarding flower scent choice, 27 bees (56.3% ± 9.5%) chose a flower associated with variable scents, and 21 bees (43.7% ± 10.8%) a flower associated with a consistent scent.

We then tested the combined effects of flower scent, colour, and random effects on bees’ first flower choice (see model (1) in Data analysis). Bees showed no preference between flowers associated with consistent or variable scents (GLMM, binomial family; intercept: Estimate = -0.36 ± 0.50, z = -0.72, *p* = 0.47; N = 48; **Fig. 2A**), and flower colour had no significant effect on this choice (X^2^ = 0.073, df = 1, *p* = 0.79).

**Figure 2:**
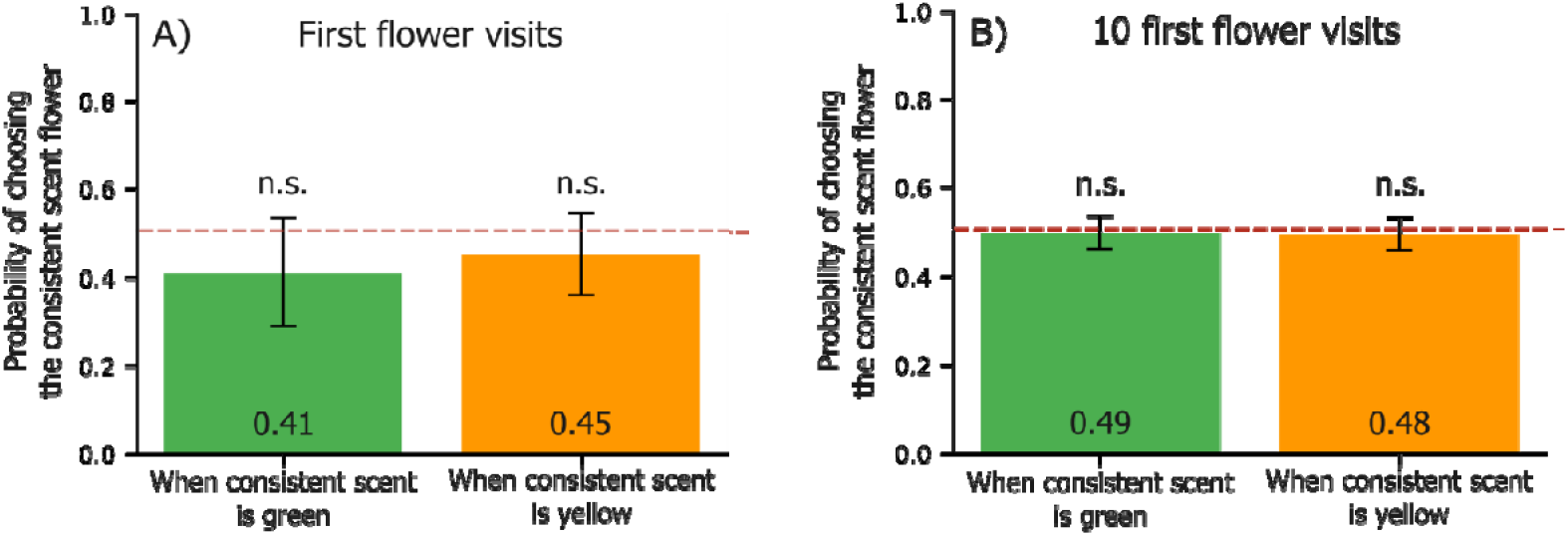
Estimated probabilities of choosing the consistent scent flower for each flower colour in the binary choice test. **A)** Proportion of choices in the first flower visit (N = 48 visits); **B)** Proportion of choices across the first 10 flower visits (N = 457 visits). Probabilities were derived from the GLMM and calculated through post hoc pairwise comparisons. The dotted red line represents a 50% chance of choosing either flower type. Error bars represent the standard errors of the predictions, and statistical differences within treatments are indicated as **n.s**. (*p* > 0.05).

### First 10 flower visits

During their first ten visits in the binary choice test, bees visited yellow flowers 51.2% (± 3.3%) of the time and green flowers 48.8% (± 3.4%). Regarding flower scent choice, 50.3% (± 3.3%) of visits were to flowers associated with variable scents, and 49.7% (± 3.3%) to flowers associated with a consistent scent.

The GLMM (see model (2) in Data Analysis) revealed no preference between flowers associated with consistent or variable scents (binomial family; intercept: Estimate = -0.007 ± 0.14, z = -0.05, *p* = 0.96; N = 457; **Fig. 2B**), and flower colour had no effect on first 10 visits (χ^2^ = 0.004, df = 1, *p* = 0.95).

### Scents preferences

Overall, bees showed no preference for any scent bouquet in the binary choice test (Chi-square goodness-of-fit test: X^2^ = 1.75, df = 5, *p* = 0.88; **Fig. 3A**) and did not favour any individual scent—lemon, rose, vanilla, or strawberry (X^2^ = 1.08, df = 3, *p* = 0.78; **Fig. 3B**).

**Figure 3:**
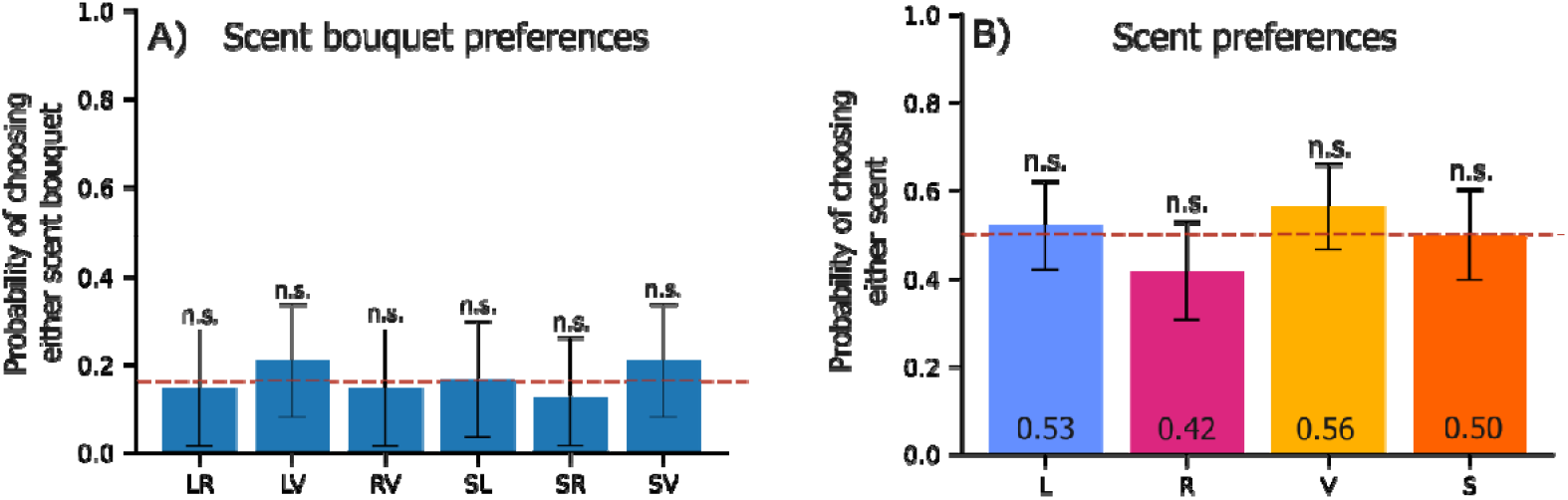
Probabilities of choosing each flower scent in the binary choice test. **(A)** Proportion of choices for each scent bouquet (N = 48 visits): lemon–rose (LR), lemon–vanilla (LV), rose–vanilla (RV), strawberry–lemon (SL), strawberry–rose (SR), or strawberry–vanilla (SV). The dotted line represents the expected random choice (16.67%). **(B)** Proportion of choices for each individual scent (N = 48 visits): Lemon (L), Rose (R), Vanilla (V), or Strawberry (S), with the dotted red line indicating a 50% chance of choosing either scent. Error bars represent standard errors, and statistical differences between scents are indicated as **n.s**. (*p* > 0.05).

## DISCUSSION

We tested whether bees had a preference between two equally rewarding flower arrays, yellow or green, one with consistent scents across flowers and the other with variables scents between flowers. Contrary to expectations, bees showed no preference for flowers with a consistent scent. During training, they readily collected sucrose solution flavoured with varying scent combinations and ratios, and in the binary choice test, they did not favour the flower colour associated with a consistent scent. To our knowledge, this is the first study to explicitly test whether bees prefer consistent or variable scents—or any other neutral cue like colour or shape—independently of reward variation.

Every tested bee collected all sucrose rewards in both flower arrays, regardless of scent composition. This reflects bees’ ability to rapidly associate scents with rewards (Kunze and Gumbert, 2001; Wright and Schiestl, 2009; Giurfa, 2007), even when they have innate preferences for specific scent volatiles (Milet-Pinheiro et al., 2013; Raguso, 2008a). Surprisingly, bees visited and accepted every flower in the variable array, despite previous findings that bumblebees exhibit even stronger flower constancy when flowers differ in multiple sensory cues, such as scents and colours (Wells and Wells, 1985; Gegear and Laverty, 2001; Gegear, 2005). However, recent research has challenged the idea of high flower constancy in bumblebees; Yourstone et al. (2023) found that bees were less flower constant than expected, with only 23% of their foraging trips occurring on the same flower species.

In the binary choice test, bees showed a slight tendency to visit yellow flowers first, though this preference was not significant. Bees are known to have an innate preference for yellow flowers (Lunau, 1990; Gumbert, 2000). To ensure that choices were based on learned associations, we used unscented, unrewarded flowers in the binary test. Hence it is unclear whether scent preferences would have outweighed colour preferences if both cues were present simultaneously. Notably, Larue et al. (2015) found that floral scent had a stronger influence than visual traits in attracting flower-visiting insects.

Were bees in our experiment able to distinguish between scent combinations, particularly the different scent blend ratios in the variable flower array? Studies show that bees can detect individual volatiles within complex scent blends, influencing their flower choices (Laloi and Pham-Delègue, 2004; Locatelli et al., 2016), and honeybees can even perceive subtle differences in the ratio of two scents (Wright et al., 2005; Bateson et al., 2011). Moreover, since our flowers contained scented sucrose solution, bees could also taste them; Robertson (2019) suggests that olfactory and gustatory receptors may overlap in bumblebees. In nature, scented nectar contains volatile organic compounds, which pollinators use to assess reward availability before landing on a flower (Heinrich, 1979; Raguso, 2008b), and can impact their foraging behaviour (Raguso, 2004). Given this, it seems likely that bees could sense the variability in the variable flowers, and distinguish this from the fixed-ratio flowers.

Although no study has directly tested bees’ preference for consistency in neutral cues, some have examined their responses to inconsistent rewarding cues. Honeybees tended to avoid choices where uncertain visual cues predict reward or punishment (Perry and Barron, 2013). Andrew et al. (2014) found that honeybees trained to discriminate between a rewarding and a punishing scent blend preferred a novel scent that was more distinct from the punishing one, rather than similar to the reward. Likewise, Lynn et al. (2005) showed that bumblebees favoured novel colours that minimised the risk of choosing an unrewarding flower. This phenomenon, known as “peak shift” (Hanson, 1959), occurs when animals develop a preference for a more extreme version of a rewarded stimulus to avoid similar, non-rewarded ones. These findings suggest that bees may prefer consistent cues over variable ones, to reduce uncertainty. Stress may further amplify this avoidance of inconsistent cues: Bateson et al. (2011) found that shaken honeybees were less likely to respond to ambiguous scents associated with rewards.

When it comes to rewards preferences, bees are generally risk-averse and tend to avoid variability (see review by Anselme, 2018). They typically prefer consistent nectar amounts over variable ones (Real, 1981; Waddington et al., 1981; Shafir et al., 1999), which is likely in part due to the psychophysics of reward perception (Kacelnik and Bateson, 1996). When resources are variable, bumblebees also forage less efficiently (Dunlap et al., 2017) and rely more on social information for flower choice (Smolla et al., 2016). However, their sensitivity to variability depends on context. In some contexts, bees showed indifference to fluctuations in nectar concentration or volume (Waddington, 1995; Fülöp and Menzel, 2000) and when variable distribution did not include null rewards (Drezner-Levy and Shafir, 2007). They may even favour variable rewards when advantageous: For example, bumblebees initially preferred consistent rewards but shifted to variable ones when colony nectar reserves were low, both in natural foraging conditions (Cartar, 1991) and in laboratory settings (Cartar and Dill, 1990).

Bees’ preferences for specific floral volatiles are well-documented (Knudsen et al., 2006; Farré-Armengol et al., 2017; Benelli et al., 2017; Bisrat and Jung, 2022), and they readily associate these scents with rewards (Wells and Wells, 1985; Wright and Schiestl, 2009; Giurfa, 2007). Bees can even retain scent memories longer than visual cues (Menzel, 1993; Kunze and Gumbert, 2001). Bumblebees, in particular, use floral scents as social cues, with foragers transferring scent compounds within the nest to inform nestmates (Molet et al., 2009). They also favour flowers that match the scents collected by successful foragers (Dornhaus and Chittka, 1999).

However, bees learn consistent scents more effectively than variable ones (Wright and Thomson, 2005; Wright and Schiestl., 2009). For instance, Wright et al. (2008) found that honeybees reject scent-modified flowers even if they contain familiar scent compounds, highlighting the importance of scent consistency for pollinator recognition. Many rewardless flowers emit weak or highly variable scent, likely to avoid detection by scent-learning pollinators (Jersáková and Johnson, 2006; Salzmann et al., 2007). Salzmann et al. (2007) found that rewarding orchids produce strong, consistent scents that bees can detect, whereas deceptive orchids emit weak, highly variable scents. Similarly, floral compounds that attract pollinators tend to be more consistent across populations and species, while non-attractive compounds show greater variability (Mant et al., 2005; Huber et al., 2005).

Nonetheless, floral scent remains a highly variable trait (Dudareva et al., 1996), even among individual flowers within the same species (Burdon et al., 2015). While plants adjust scent emissions to attract pollinators (Majetic et al., 2015; Burkle et al., 2020), scent bouquets are also influenced by environmental factors (Dudareva et al., 1999; Raguso, 2008b). Studies suggest that intraspecific variation in floral scent may help attract local pollinators (Soler et al., 2011; Larue et al., 2016; Vega-Polanco et al., 2023). Fülöp and Menzel (2000) propose that bees’ tolerance for scent variability may be an adaptation to cope with unreliable floral resources.

Floral scents, including scented nectar, likely serve as honest signals to pollinators (Howell and Alarcón, 2007; Gervasi and Schiestl, 2017). Bees often select flowers based on their scent composition (Pichersky and Raguso, 2018; Knudsen and Gershenzon, 2020) or the intensity of scent emissions (Majetic et al., 2009; Parachnowitsch et al., 2013), which inform foragers about nectar and pollen availability. We suggest that bees are likely to respond to olfactory cues regardless of whether they are more variable or consistent, as long as they can reliably associate them with rewards.

Our findings highlight bees’ ability to rapidly learn scent-reward associations and show that they forage equally on flower arrays with both variable and consistent scents, and show no preference between them. This in turn suggests that plants may not be under strong selection to reduce inter-flower scent variability, at least in terms compound ratios. Maintaining tight control over a trait in the face of environmental variation is costly, and if no selection for this maintenance is present, we would expect such traits to fluctuate, as indeed is the case with intraspecific scent variation between flowers (Burdon et al., 2015; Delle-Vedove et al., 2017).

However, our controlled experiment likely oversimplified the complex sensory landscape bees navigate in nature, and further research is needed to understand how they respond to scent variability within flower patches. Despite its importance, the role of floral scent in bee foraging has been largely overlooked (Fenster et al., 2004; Raguso, 2004, 2008). More generally, future studies should examine whether bees’ aversion to variability in rewards extends to other neutral cues like morphology or colour, as the foraging preferences of plant pollinators play a key role in shaping floral trait evolution (Ollerton et al., 2011).

## SUPPLEMENTARY MATERIAL

**S1** contains the dataset of the main experiment, **S2** provides the reflectance curves of the artificial flowers, and **S3** the statistical analysis of the experiment. All supplementary materials are available on Zenodo (https://doi.org/10.5281/zenodo.14993082).

## ACKNOWLEDGMENTS

We want to thank A. Koch and M. Kietniz for their assistance with data collection, and K. Hartmannsgruber for video analysis. Special thanks to A. Avarguès-Weber for providing spectrometer measurements of the artificial flowers.

## AUTHORS’ CONTRIBUTION

**Mélissa Armand:** Conceptualization, Methodology, Software, Validation, Formal analysis, Investigation, Writing -Original Draft, Writing -Review and Editing, Visualization. **Lisa Zeilmann:** Investigation. **Christian Weinzettl**: Investigation. **Tomer J. Czaczkes:** Conceptualization, Methodology, Validation, Resources, Writing -Review and Editing, Supervision, Project administration, Funding acquisition.

## REFERENCES

Andrew SC, Perry CJ, Barron AB, et al (2014) Peak shift in honey bee olfactory learning. Anim Cogn 17:1177–1186. 10.1007/s10071-014-0750-3

Anselme P (2018) Uncertainty processing in bees exposed to free choices: Lessons from vertebrates. Psychon Bull Rev 25:2024–2036. 10.3758/s13423-018-1441-x

Armengol GF Biotic and abiotic factors that determine the emission of volatile organic compounds by flowers Bateson M, Desire S, Gartside SE, Wright GA (2011) Agitated Honeybees Exhibit Pessimistic Cognitive Biases. Current Biology 21:1070–1073. 10.1016/j.cub.2011.05.017

Benelli G, Canale A, Romano D, et al (2017) Flower scent bouquet variation and bee pollinator visits in Stevia rebaudiana Bertoni (Asteraceae), a source of natural sweeteners. Arthropod-Plant Interactions 11:381–388. 10.1007/s11829-016-9488-y

Bisrat D, Jung C (2022) Roles of flower scent in bee-flower mediations: a review. Journal of Ecology and Environment 46:18–30. 10.5141/jee.21.00075

Brooks ME, Kristensen K, van Benthem KJ, et al (2017) glmmTMB balances speed and flexibility among packages for zero-inflated generalized linear mixed modeling. The R journal 9:378–400. 10.3929/ethz-b-000240890

Burdon RCF, Raguso RA, Kessler A, Parachnowitsch AL (2015) Spatiotemporal Floral Scent Variation of Penstemon digitalis. J Chem Ecol 41:641–650. 10.1007/s10886-015-0599-1

Burkle LA, Glenny WR, Runyon JB (2020) Intraspecific and interspecific variation in floral volatiles over time. Plant Ecol 221:529–544. 10.1007/s11258-020-01032-1

Cartar RV (1991) A Test of Risk-Sensitive Foraging in Wild Bumble Bees. Ecology 72:888–895. 10.2307/1940590

Cartar RV, work(s): LMDR (1990) Colony Energy Requirements Affect the Foraging Currency of Bumble Bees. Behavioral Ecology and Sociobiology 27:377–383

Chittka L, Thomson JD, Waser NM (1999) Flower Constancy, Insect Psychology, and Plant Evolution. Naturwissenschaften 86:361–377. 10.1007/s001140050636

Delle-Vedove R, Schatz B, Dufay M (2017) Understanding intraspecific variation of floral scent in light of evolutionary ecology. Annals of Botany 120:1–20. 10.1093/aob/mcx055

Dobson HEM (1994) Floral Volatiles in Insect Biology. In: Insect-Plant Interactions (1993). CRC Press

Dobson HEM (ed) (2006) Relationship between Floral Fragrance Composition and Type of Pollinator. In: Biology of Floral Scent. CRC Press

Dornhaus A, Chittka L (1999) Evolutionary origins of bee dances. Nature 401:38–38. 10.1038/43372

Drezner-Levy T, Shafir S (2007) Parameters of variable reward distributions that affect risk sensitivity of honey bees. Journal of Experimental Biology 210:269–277. 10.1242/jeb.02656

Dudareva N, Cseke L, Blanc VM, Pichersky E (1996) Evolution of floral scent in Clarkia: novel patterns of S-linalool synthase gene expression in the C. breweri flower. The Plant Cell 8:1137–1148. 10.1105/tpc.8.7.1137

Dudareva N, Pichersky E, Gershenzon J (2004) Biochemistry of Plant Volatiles. Plant Physiology 135:1893–1902. 10.1104/pp.104.049981

Dunlap AS, Papaj DR, Dornhaus A (2017) Sampling and tracking a changing environment: persistence and reward in the foraging decisions of bumblebees. Interface Focus 7:20160149. 10.1098/rsfs.2016.0149

Farré-Armengol G, Fernández-Martínez M, Filella I, et al (2020) Deciphering the Biotic and Climatic Factors That Influence Floral Scents: A Systematic Review of Floral Volatile Emissions. Front Plant Sci 11:1154. 10.3389/fpls.2020.01154

Farré-Armengol G, Filella I, Llusià J, Peñuelas J (2017) ?-Ocimene, a Key Floral and Foliar Volatile Involved in Multiple Interactions between Plants and Other Organisms. Molecules 22:1148. 10.3390/molecules22071148

Fenster CB, Armbruster WS, Wilson P, et al (2004) Pollination Syndromes and Floral Specialization. Annual Review of Ecology, Evolution, and Systematics 35:375–403. 10.1146/annurev.ecolsys.34.011802.132347

Fülöp A, Menzel R (2000) Risk-indifferent foraging behaviour in honeybees. Animal Behaviour 60:657–666. 10.1006/anbe.2000.1492

Gegear RJ (2005) Multicomponent floral signals elicit selective foraging in bumblebees. Naturwissenschaften 92:269–271. 10.1007/s00114-005-0621-5

Gegear RJ, Laverty TM (2005) Flower constancy in bumblebees: a test of the trait variability hypothesis. Animal Behaviour 69:939–949. 10.1016/j.anbehav.2004.06.029

Gervasi DDL, Schiestl FP (2017a) Real-time divergent evolution in plants driven by pollinators. Nat Commun 8:14691. 10.1038/ncomms14691

Giurfa M (2007) Behavioral and neural analysis of associative learning in the honeybee: a taste from the magic well. J Comp Physiol A 193:801–824. 10.1007/s00359-007-0235-9

Goulson D (1999) Foraging strategies of insects for gathering nectar and pollen, and implications for plant ecology and evolution. Perspectives in Plant Ecology, Evolution and Systematics 2:185–209. 10.1078/1433-8319-00070

Goulson D, Stout JC, Langley J, Hughes WOH (2000) Identity and Function of Scent Marks Deposited by Foraging Bumblebees. J Chem Ecol 26:2897–2911. 10.1023/A:1026406330348

Gumbert A (2000) Color choices by bumble bees (Bombus terrestris): innate preferences and generalization after learning. Behav Ecol Sociobiol 48:36–43. 10.1007/s002650000213

Hanson HM (1959) Effects of discrimination training on stimulus generalization. Journal of Experimental Psychology 58:321–334. 10.1037/h0042606

Hartig F (2020) DHARMa: residual diagnostics for hierarchical (multi-level/mixed) regression models. R package version 03 3:

Heinrich B (1977) Pollination Energetics: An Ecosystem Approach. In: Mattson WJ (ed) The Role of Arthropods in Forest Ecosystems. Springer, Berlin, Heidelberg, pp 41–46

Heinrich B (1979) Resource heterogeneity and patterns of movement in foraging bumblebees. Oecologia 40:235–245. 10.1007/BF00345321

Howell AD, Alarcón R (2007) Osmia bees (Hymenoptera: Megachilidae) can detect nectar-rewarding flowers using olfactory cues. Animal Behaviour 74:199–205. 10.1016/j.anbehav.2006.11.012

Huber FK, Kaiser R, Sauter W, Schiestl FP (2005) Floral scent emission and pollinator attraction in two species of Gymnadenia (Orchidaceae). Oecologia 142:564–575. 10.1007/s00442-004-1750-9

Hunter JD (2007) Matplotlib: A 2D Graphics Environment. Computing in Science & Engineering 3:90–95. 10.1109/MCSE.2007.55

Jersáková J, Johnson SD (2006) Lack of floral nectar reduces self-pollination in a fly-pollinated orchid. Oecologia 147:60–68. 10.1007/s00442-005-0254-6

Kacelnik A, Bateson M (1996) Risky Theories—The Effects of Variance on Foraging Decisions1. American Zoologist 36:402–434. 10.1093/icb/36.4.402

Kevan PG, Baker HG (1983) Insects as Flower Visitors and Pollinators. Annu Rev Entomol 28:407–453. 10.1146/annurev.en.28.010183.002203

Klein A-M, Vaissière BE, Cane JH, et al (2006) Importance of pollinators in changing landscapes for world crops. Proceedings of the Royal Society B: Biological Sciences 274:303–313. 10.1098/rspb.2006.3721

Knudsen JT, Eriksson R, Gershenzon J, Ståhl B (2006) Diversity and distribution of floral scent. Bot Rev 72:1–120. 10.1663/0006-8101(2006)72[1:DADOFS]2.0.CO;2

Knudsen JT, Gershenzon J (2020) The Chemical Diversity of Floral Scent. In: Biology of Plant Volatiles, 2nd edn. CRC Press

Kunze J, Gumbert A (2001) The combined effect of color and odor on flower choice behavior of bumble bees in flower mimicry systems. Behavioral Ecology 12:447–456. 10.1093/beheco/12.4.447

Laloi D, Pham-Delègue M-H (2004) Bumble Bees Show Asymmetrical Discrimination Between Two Odors in a Classical Conditioning Procedure. Journal of Insect Behavior 17:385–396. 10.1023/B:JOIR.0000031538.15346.e1

Larue A-AC, Raguso RA, Junker RR (2016) Experimental manipulation of floral scent bouquets restructures flower–visitor interactions in the field. Journal of Animal Ecology 85:396–408. 10.1111/1365-2656.12441

Lenth RV (2020) Emmeans: Estimated Marginal Means, Aka Least-Squares Means. R Package Version 1.5. 3. CRAN

Leonard AS, Dornhaus A, Papaj DR (2011) Flowers help bees cope with uncertainty: signal detection and the function of floral complexity. Journal of Experimental Biology 214:113–121. 10.1242/jeb.047407

Levin RA, McDade LA, Raguso RA (2003) The Systematic Utility of Floral and Vegetative Fragrance in Two Genera of Nyctaginaceae. Systematic Biology 52:334–351. 10.1080/10635150390196975

Locatelli FF, Fernandez PC, Smith BH (2016) Learning about natural variation of odor mixtures enhances categorization in early olfactory processing. Journal of Experimental Biology 219:2752–2762. 10.1242/jeb.141465

Loughrin JN, Hamilton-Kemp TR, Andersen RA, Hildebrand DF (1990) Volatiles from flowers of Nicotiana sylvestris, N. otophora and Malus × domestica: headspace components and day/night changes in their relative concentrations. Phytochemistry 29:2473–2477. 10.1016/0031-9422(90)85169-G

Lunau K (1990) Colour saturation triggers innate reactions to flower signals: Flower dummy experiments with bumblebees. J Comp Physiol A 166:827–834. 10.1007/BF00187329

Lynn SK, Cnaani J, Papaj DR (2005) Peak Shift Discrimination Learning as a Mechanism of Signal Evolution. Evolution 59:1300–1305. 10.1111/j.0014-3820.2005.tb01780.x

Majetic CJ, Raguso RA, Ashman T-L (2009) The sweet smell of success: floral scent affects pollinator attraction and seed fitness in Hesperis matronalis. Functional Ecology 23:480–487. 10.1111/j.1365-2435.2008.01517.x

Mant J, Peakall R, Schiestl FP (2005) Does Selection on Floral Odor Promote Differentiation Among Populations and Species of the Sexually Deceptive Orchid Genus Ophrys? Evolution 59:1449–1463. 10.1111/j.0014-3820.2005.tb01795.x

McKinney W (2010) Data Structures for Statistical Computing in Python. Austin, Texas, pp 56–61

Menzel R (1993) Associative learning in honey bees. Apidologie 24:157–168. 10.1051/apido:19930301

Milet-Pinheiro P, Ayasse M, Dobson HE, et al (2013) The chemical basis of host-plant recognition in a specialized bee pollinator. Journal of chemical ecology 39:1347–1360

Molet M, Chittka L, Raine NE (2009) How floral odours are learned inside the bumblebee (Bombus terrestris) nest. Naturwissenschaften 96:213–219. 10.1007/s00114-008-0465-x

Ollerton J, Winfree R, Tarrant S (2011) How many flowering plants are pollinated by animals? Oikos 120:321–326. 10.1111/j.1600-0706.2010.18644.x

Parachnowitsch A, Burdon RCF, A. Raguso R, Kessler A (2013) Natural selection on floral volatile production in Penstemon digitalis: Highlighting the role of linalool. Plant Signaling & Behavior 8:e22704. 10.4161/psb.22704

Perry CJ, Barron AB (2013) Honey bees selectively avoid difficult choices. Proceedings of the National Academy of Sciences 110:19155–19159. 10.1073/pnas.1314571110

Pichersky E, Raguso RA (2018) Why do plants produce so many terpenoid compounds? New Phytologist 220:692–702. 10.1111/nph.14178

Pichersky E, Raguso RA, Lewinsohn E, Croteau R (1994) Floral Scent Production in Clarkia (Onagraceae) (I. Localization and Developmental Modulation of Monoterpene Emission and Linalool Synthase Activity). Plant Physiology 106:1533–1540. 10.1104/pp.106.4.1533

Raguso RA (2004) Why Are Some Floral Nectars Scented? Ecology 85:1486–1494. 10.1890/03-0410

Raguso RA (2008a) Start making scents: the challenge of integrating chemistry into pollination ecology. Entomologia Experimentalis et Applicata 128:196–207. 10.1111/j.1570-7458.2008.00683.x

Raguso RA (2008b) Wake Up and Smell the Roses: The Ecology and Evolution of Floral Scent. Annual Review of Ecology, Evolution, and Systematics 39:549–569. 10.1146/annurev.ecolsys.38.091206.095601

Raguso RA (2020) Behavioral Responses to Floral Scent: Experimental Manipulations and Multimodal Plant–Pollinator Communication. In: Biology of Plant Volatiles, 2nd edn. CRC Press

Raguso RA, Levin RA, Foose SE, et al (2003) Fragrance chemistry, nocturnal rhythms and pollination “syndromes” in Nicotiana. Phytochemistry 63:265–284. 10.1016/S0031-9422(03)00113-4

Raguso RA, Pichersky E (1999) New Perspectives in Pollination Biology: Floral Fragrances. A day in the life of a linalool molecule: Chemical communication in a plant-pollinator system. Part 1: Linalool biosynthesis in flowering plants. Plant Species Biology 14:95–120. 10.1046/j.1442-1984.1999.00014.x

Raine NE, Chittka L (2008) The correlation of learning speed and natural foraging success in bumble-bees. Proceedings of the Royal Society B: Biological Sciences 275:803–808. 10.1098/rspb.2007.1652

Raine NE, Chittka L (2007) The Adaptive Significance of Sensory Bias in a Foraging Context: Floral Colour Preferences in the Bumblebee Bombus terrestris. PLOS ONE 2:e556. 10.1371/journal.pone.0000556

Real LA (1981) Uncertainty and Pollinator-Plant Interactions: The Foraging Behavior of Bees and Wasps on Artificial Flowers. Ecology 62:20–26. 10.2307/1936663

Robertson HM (2019) Molecular Evolution of the Major Arthropod Chemoreceptor Gene Families. Annual Review of Entomology 64:227–242. 10.1146/annurev-ento-020117-043322

Saleh N, Scott AG, Bryning GP, Chittka L (2007) Distinguishing signals and cues: bumblebees use general footprints to generate adaptive behaviour at flowers and nest. Arthropod-Plant Interactions 1:119–127. 10.1007/s11829-007-9011-6

Salzmann CC, Nardella AM, Cozzolino S, Schiestl FP (2007) Variability in Floral Scent in Rewarding and Deceptive Orchids: The Signature of Pollinator-imposed Selection? Annals of Botany 100:757–765. 10.1093/aob/mcm161

Schiestl FP, Dötterl S (2012) The evolution of floral scent and olfactory preferences in pollinators: Coevolution or pre-existing bias? Evolution 66:2042–2055. 10.1111/j.1558-5646.2012.01593.x

Schiestl FP, Johnson SD (2013) Pollinator-mediated evolution of floral signals. Trends in Ecology & Evolution 28:307–315. 10.1016/j.tree.2013.01.019

Shafir S, Wiegmann DD, Smith BH, Real LA (1999) Risk-sensitive foraging: choice behaviour of honeybees in response to variability in volume of reward. Animal Behaviour 57:1055–1061. 10.1006/anbe.1998.1078

Smolla M, Alem S, Chittka L, Shultz S (2016) Copy-when-uncertain: bumblebees rely on social information when rewards are highly variable. Biology Letters 12:20160188. 10.1098/rsbl.2016.0188

Soler C, Hossaert-McKey M, Buatois B, et al (2011) Geographic variation of floral scent in a highly specialized pollination mutualism. Phytochemistry 72:74–81. 10.1016/j.phytochem.2010.10.012

Solís-Montero L, Cáceres-García S, Alavez-Rosas D, et al (2018) Pollinator Preferences for Floral Volatiles Emitted by Dimorphic Anthers of a Buzz-Pollinated Herb. J Chem Ecol 44:1058–1067. 10.1007/s10886-018-1014-5

Sprayberry JDH (2018) The prevalence of olfactory-versus visual-signal encounter by searching bumblebees. Sci Rep 8:14590. 10.1038/s41598-018-32897-y

Tan KH, Nishida R (2012) Methyl eugenol: Its occurrence, distribution, and role in nature, especially in relation to insect behavior and pollination. Journal of Insect Science 12:56. 10.1673/031.012.5601

Vega-Polanco M, Solís-Montero L, Rojas JC, et al (2023) Intraspecific variation of scent and its impact on pollinators’ preferences. AoB PLANTS 15:plad049. 10.1093/aobpla/plad049

Waddington KD (1995) Bumblebees Do Not Respond to Variance in Nectar Concentration. Ethology 101:33–38. 10.1111/j.1439-0310.1995.tb00342.x

Waddington KD, Allen T, Heinrich B (1981) Floral preferences of bumblebees (Bombus edwardsii) in relation to intermittent versus continuous rewards. Animal Behaviour 29:779–784. 10.1016/S0003-3472(81)80011-5

Waser NM (1986) Flower Constancy: Definition, Cause, and Measurement. The American Naturalist 127:593–603. 10.1086/284507

Wells PH, Wells H (1985) Ethological Isolation of Plants 2. Odour Selection By Honeybees. Journal of Apicultural Research 24:86–92. 10.1080/00218839.1985.11100654

Whibley AC, Langlade NB, Andalo C, et al (2006) Evolutionary Paths Underlying Flower Color Variation in Antirrhinum. Science 313:963–966. 10.1126/science.1129161

Wright GA, Kottcamp SM, Thomson MGA (2008) Generalization Mediates Sensitivity to Complex Odor Features in the Honeybee. PLOS ONE 3:e1704. 10.1371/journal.pone.0001704

Wright GA, Lutmerding A, Dudareva N, Smith BH (2005) Intensity and the ratios of compounds in the scent of snapdragon flowers affect scent discrimination by honeybees (Apis mellifera). J Comp Physiol A 191:105–114. 10.1007/s00359-004-0576-6

Wright GA, Schiestl FP (2009) The evolution of floral scent: the influence of olfactory learning by insect pollinators on the honest signalling of floral rewards. Functional Ecology 23:841–851. 10.1111/j.1365-2435.2009.01627.x

Wright GA, Thomson MGA (2005) Odor Perception and the Variability in Natural Odor Scenes. In: Recent Advances in Phytochemistry. Elsevier, pp 191–226

Yourstone J, Varadarajan V, Olsson O (2023) Bumblebee flower constancy and pollen diversity over time. Behavioral Ecology 34:602–612. 10.1093/beheco/arad028

